# Lysine as a potential low molecular weight angiogen: its clinical, experimental and in-silico validation- A brief study

**DOI:** 10.1101/080176

**Authors:** Debatosh Datta, Priyanshu Verma, Anindita Banerjee, Sujoy Kar, Tanima Sengupta, Nalinava Sengupta, Sujoy Kumar Samanta, Enam Murshed Khan

## Abstract

Globally, the area of angiogenesis is dominated by investigations on anti-angiogenic agents and processes, due to its role in metastatic cancer treatment. Although, the area of ischemic tissue reperfusion is having much bigger demand and foot-mark. Following clinical failure of VEGF (Vascular endothelial growth factor) as a potential agent for induction of a controlled angiogenic response in ischemic tissues and organs, the progress is reasonably quiet as for new low molecular weight (LMW) angiogen molecules and their clinical applications are concerned. Basic amino acid Lysine has been observed to have profound angiogenic property in ischemic tissues, which is controlled, reproducible, time bound and without any accompanying reperfusion damage. In this study, the basic amino acid Lysine has been suggested as a LMW-angiogen, where it has been proposed to have a molecular binding property between VEGF and VEGF receptor (VEGFR). Here, the molecular adhesive hypothesis is being probed and confirmed both in the clinical and lab conditions through induced angiogenic response in tissue repair and in chick chorio allantoic membrane (CAM), respectively; and in dry-docking experiments (in-silico studies).

## 1. Introduction

Controlled reperfusion of ischemic tissues is an active area of investigation both as an alternate approach of therapy and as the fundamental platform of in-situ cellular expansion in healing of wounds and restoration of blood supply in ischemic tissues. VEGF is the lead mitogen for endothelial cells. In human subjects, VEGF family consist of five different isoforms - VEGF-A, VEGF-B, VEGF-C, VEGF-D and PlGF (placental growth factor)^1^. It plays a central role in the initiation of angiogenesis: regeneration and growth of new blood vessels at the capillary level. In vascular exchange beds, angiogenesis, which is an extremely conserved process across species, forms network of the supply lines for healthy cells by providing them required nutrients and gas, signaling molecules and by removal of carbon dioxide and other metabolic end products^1-6^. As mentioned, angiogenesis is a highly conserved natural process and endogenously controlled by the body’s own regulatory system. This system is considerably complex and precise, which involve both endogenous inhibitors and activators^6^. Degree of angiogenesis supports natural growth phase in the early part of life. However, with aging “normal angiogenic process” proves slow and inadequate for required growth at any given instance and in situations of very high demand for tissue repair and significant tissue regeneration. In-vivo angiogenesis is promoted through the production of VEGF by endothelial cells in the domains of relative ischaemia followed by its binding to the dedicated receptor VEGFR in the immediate vicinity. In addition, the VEGFR family consists of two non-protein kinases (neuropilin-1 and 2) and three receptor tyrosine kinases (VEGFR-1, VEGFR-2 and VEGFR-3) ^1,4-6^. VEGFR-1 binds to VEGF-A, VEGF-B and PlGF; VEGFR-2 binds to VEGF-A, VEGF-C and VEGF-D; VEGFR-3 binds to VEGF-C and VEGF-D. VEGFR-1 and VEGFR-2 receptors are mainly found on the blood vascular endothelium and responsible for angiogenesis. However, VEGFR-3 receptors are mainly found on the lymphatic endothelium and responsible for lymphangiogenesis^2-4^. Moreover, in situations of relative ischaemia, depending on the degree, ischaemic zone endothelial cells elaborate excess titres of VEGF as a means of controlled ischaemia reperfusion^7,8^.

Since, tumor angiogenesis or uncontrolled non-hierarchical angiogenesis play central role in proliferation of cancerous cells, the area of angiogenesis is mainly governed by studies on the inhibition of the process^9-14^. In addition, the absence of a good candidate molecule as an effective mediator of induced angiogenic process the field of induced controlled augmented angiogenesis is substantially on a low key in the current global research trend.

Following clinical limitation of use of VEGF as a therapeutic agent, the area of controlled angiogenesis is evolving rather slowly, because of the absence of a non-toxic low-molecular-weight angiogen molecule compared to rapid progresses made in the other half of angiogenesis research i.e. suppression of it^15^.

In this study, the potential of basic amino acid Lysine as a LMW-angiogen has been examined with the help of histopathological and CAM (Chorio-Allantoic-Membrane) experiments. Molecular docking or in-silico docking has been employed to check and compare the binding affinities of VEGF-A and VEGF-A receptor with Lysine. In addition, experimentally observed results have been further validated through the in-silico docking studies of VEGF-A and VEGF-A receptor in the presence of Lysine.

## 2. Materials and Methods

### 2.1. Histopathological assay

Histopathological assay involved a surface wound treatment with bare 24-hourly topical application of hydrogel based Lysine HCl formulation (250 mg/g) to observe inducible angiogenesis in wound bed, if any. Human study was undertaken under all prescribed rules governing clinical evaluation studies, both at the controller general of drugs as well as the local institutional ethical committee levels. In addition, informed and explained patient consent was part of this clinical study. Following topical application for 5 days under moistened wound conditions, routine histopathology studies were carried out with tissue samples obtained from the peripheral margin of wound beds through punch biopsy.

### 2.2. CAM experiment

Chorio-Allantoic-Membrane (CAM) model was utilized to have confirmatory visual evidence of inducible angiogenesis as described earlier^14,16-18^. Briefly, CAM of 9-days-old chicken embryos had been used in two groups: experimental group versus a control group, each group contains 5 embryos. Control CAMs were installed with aqueous vehicle (2 ml of water) only, whereas experimental group were supplemented with 2 ml of an aqueous solution of Lysine HCl (Ajinomoto Co., Japan). Both the sets of embryos were photographed after the 5 days incubation at 37 °C (to visualize the sprouting vessels).

### 2.3. In-silico docking analysis

The present section is mainly focused to the in-silico study of binding affinity of VEGF-A to VEGF-A receptor in the presence of basic amino acid Lysine. Docking tools -Z-DOCK^19^ and PATCHDOCK^20^ followed by FIREDOCK^21,22^ have been used for the analyses. Required protein molecules’ pdb files have been selected from the PDB database^23^. Molecules with PDB IDs -1VPF (VEGF-A) and 4KZN (VEGF-A receptor) have been used respectively for the docking studies. The Lysine molecule’s sdf file has been downloaded from the PubChem server^24^ with further conversion to pdb file with the help of an offline tool - Open Babel^25^.

Prediction studies have been performed for the same combination to predict the most probable binding site(s) on Z-DOCK platform. However, for stability analysis, only FIREDOCK results have been used. Chimera 1.10.2^26^ and Swiss-PdbViewer 4.1.0^27^ has been used for the visualization of models generated through different docking tools.

This study is mainly based on a hypothesis – basic amino acid Lysine possibly acts as a molecular binder between the most potent in-vivo angiogenic peptide VEGF and its receptor, thereby augmenting the binding between the two resulting in an enhanced biological response (Fig. 1)^28^.

**Fig. 1.**
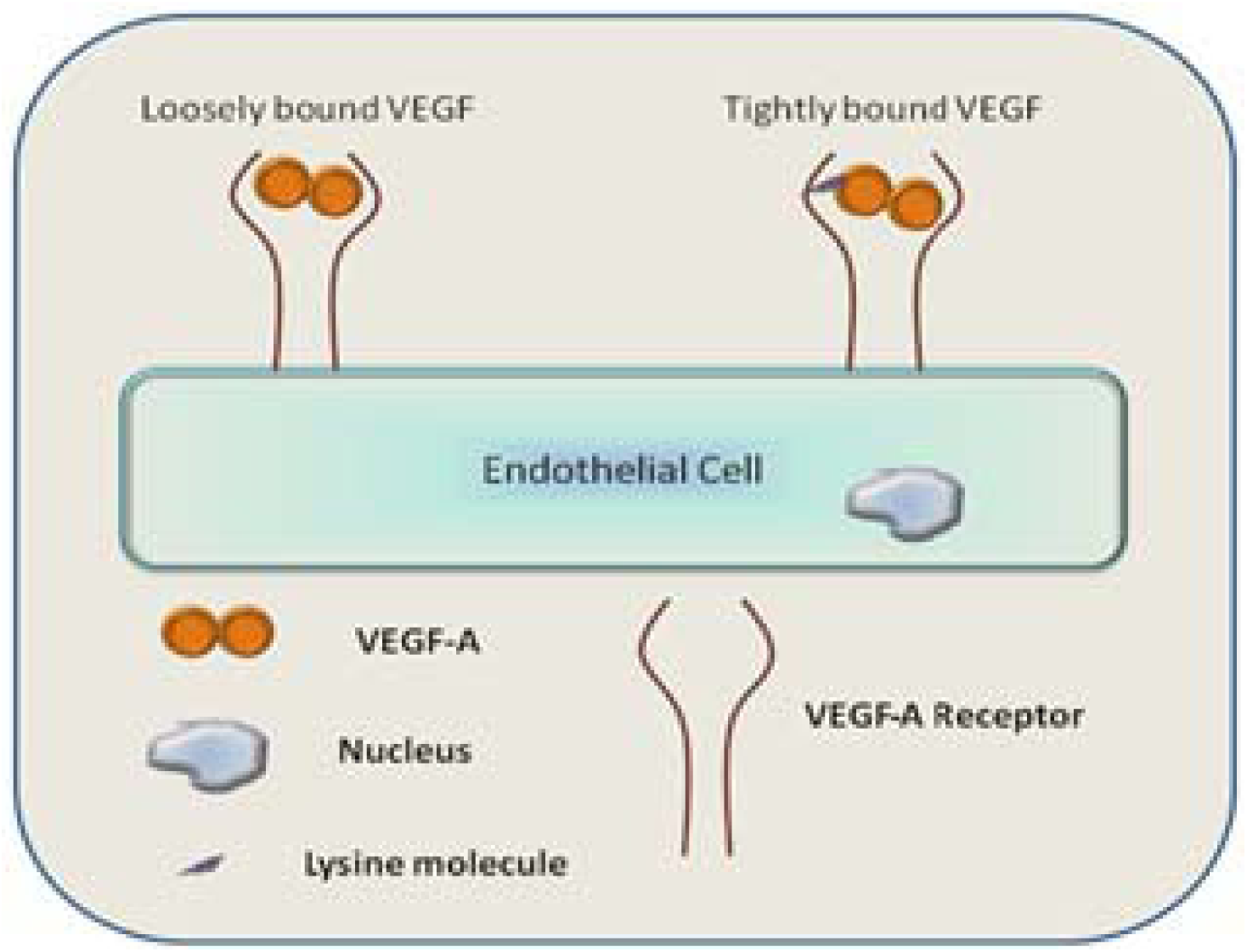
Schematic representation of the hypothesis: VEGF-A and VEGF-A receptor binding on the endothelial cell surface in the presence of Lysine molecule.

This facilitated binding probably enhances the in-vivo physiological level of angiogenic process which otherwise helps in controlled reperfusion of ischaemic tissues anywhere despite being remarkably slow.

To examine the hypothesis, the present design included two modes of approach in in-silico binding studies:

i. Lysine being coupled to VEGF-A receptor (4KZN) to generate the complex–VEGF-A receptor-Lysine. This complex is then presented for binding with VEGF-A (1VPF).
ii. Lysine is first coupled to VEGF-A (1VPF) to generate the complex–VEGF-A-Lysine. This complex peptide is then presented for binding studies with VEGF-A receptor (4KZN).

In protocol (i) above, VEGF-A receptor structure was scanned for distribution of surface negative charges along the entire length of the peptide. This was done in anticipation of a possible electrostatic interaction between added Lysine and the receptor peptide (Fig. 2a). Similarly, in pre-binding added lysine complexing protocol (ii) described above, VEGF-A molecule was screened for negative charge distribution along the entire length of the peptide molecule (Fig. 2d).

**Fig. 2.**
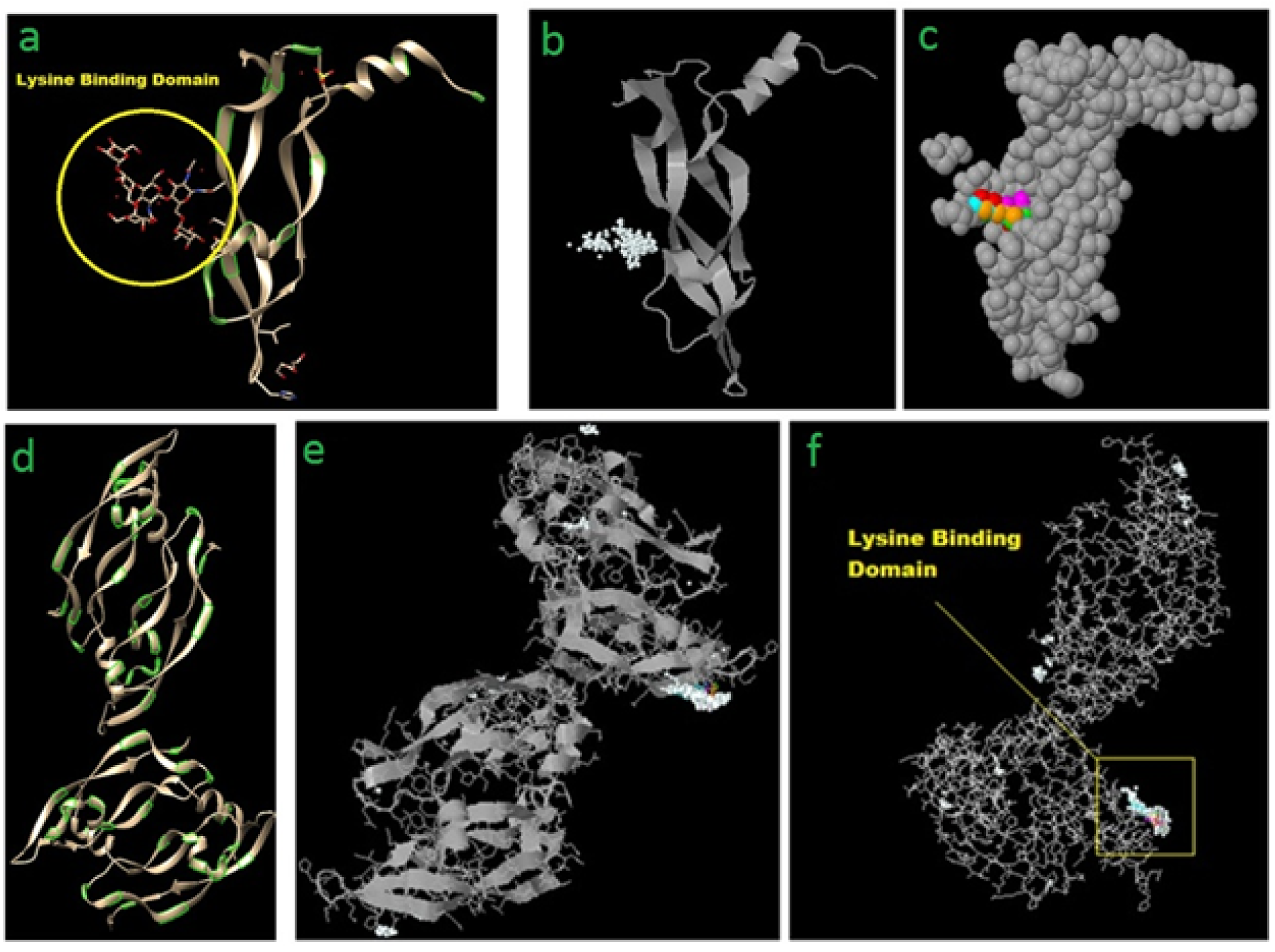
**(a)** VEGF-A receptor (4KZN) showing the most probable lysine binding domain as well as negatively charged amino acid(s) in the vicinity (mainly aspartic acid) marked as green islands. Z-DOCK results for 4KZN (VEGF-A receptor) and Lysine (as a ligand) showing top 500 possible Lysine binding sites **(b)**, and top 5 Lysine binding sites **(c)** on VEGF-A receptor. **(d)** VEGF-A (1VPF) ribbon structure showing uniform distribution of negatively charged amino acids along the entire length of the chains. Z-DOCK results for 1VPF (VEGF-A) and Lysine (as an additional ligand) showing ribbon structure **(e)**, and wire-frame structure **(f)** of VEGF-A (1VPF) with top 500 preferential binding locations of added Lysine residue (white spheres) and top 5 locations (colored lines).

## 3. Results and Discussions

### 3.1. Histopathological assay

Lysine induced remarkably enhanced angiogenic response in the surface tissue repair of extensive bedsore was observed within approx. two weeks’ time (Fig.3a). Low and high power field optical microscopic representations of the tissue obtained from the margin of the wound bed through punch biopsy showed hierarchical angiogenic response in the wound bed (Fig. 3b, c). This clinical evidence of topical Lysine HCl gel mediated induction of an extensive angiogenic response in a profusely ischaemic wound bed possibly conforms to the hypothesis of the enhanced angiogenic response capability of the natural amino acid Lysine in ischemic tissues.

**Fig. 3.**
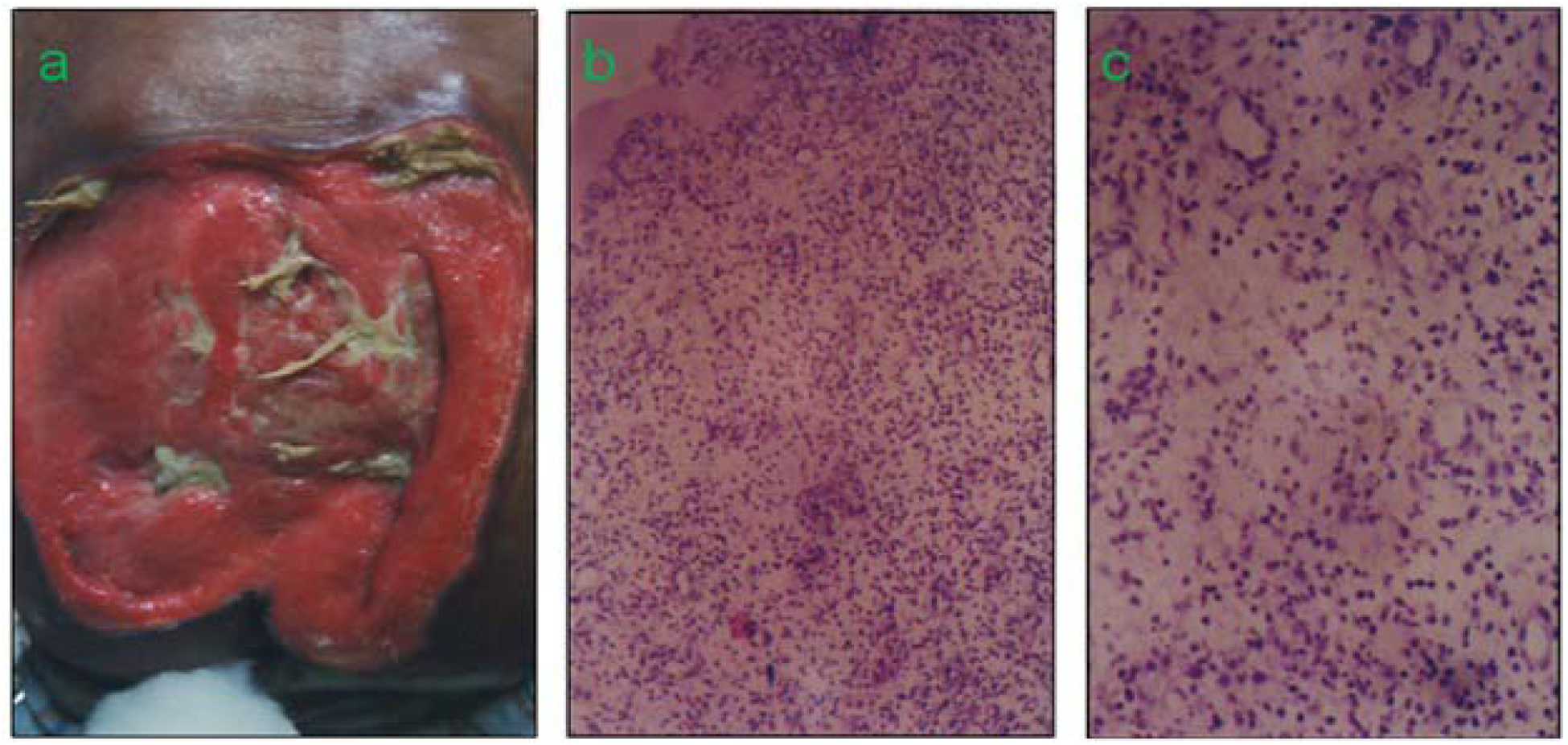
**(a)** Extensive bedsore in a clinical study (patient was ready for graft with two-weeks’ topical application of a Lysine hydrogel formulation 250 mg/g). **(b)** Low-power field optical microscopic representation of the tissue from the margin of the wound obtained through punch biopsy. **(c)** High power field photograph of the tissue showing the hierarchical angiogenic response in the wound bed.

### 3.2. CAM experiment

The ability of the amino acid as a potent angiogen was evaluated in in-vitro angiogenesis model. Chicken embryo CAMs were supplemented with Lysine HCl in aqueous form (250 mg/ml). A representative photograph showed sharp quantitative contrast in the angiogenic response in chicken embryo CAMs with and without Lysine in terms of growth of new capillaries (Fig. 4a, b). Substantive difference between the natural and lysine induced angiogenic response is obvious through the figures.

**Fig. 4.**
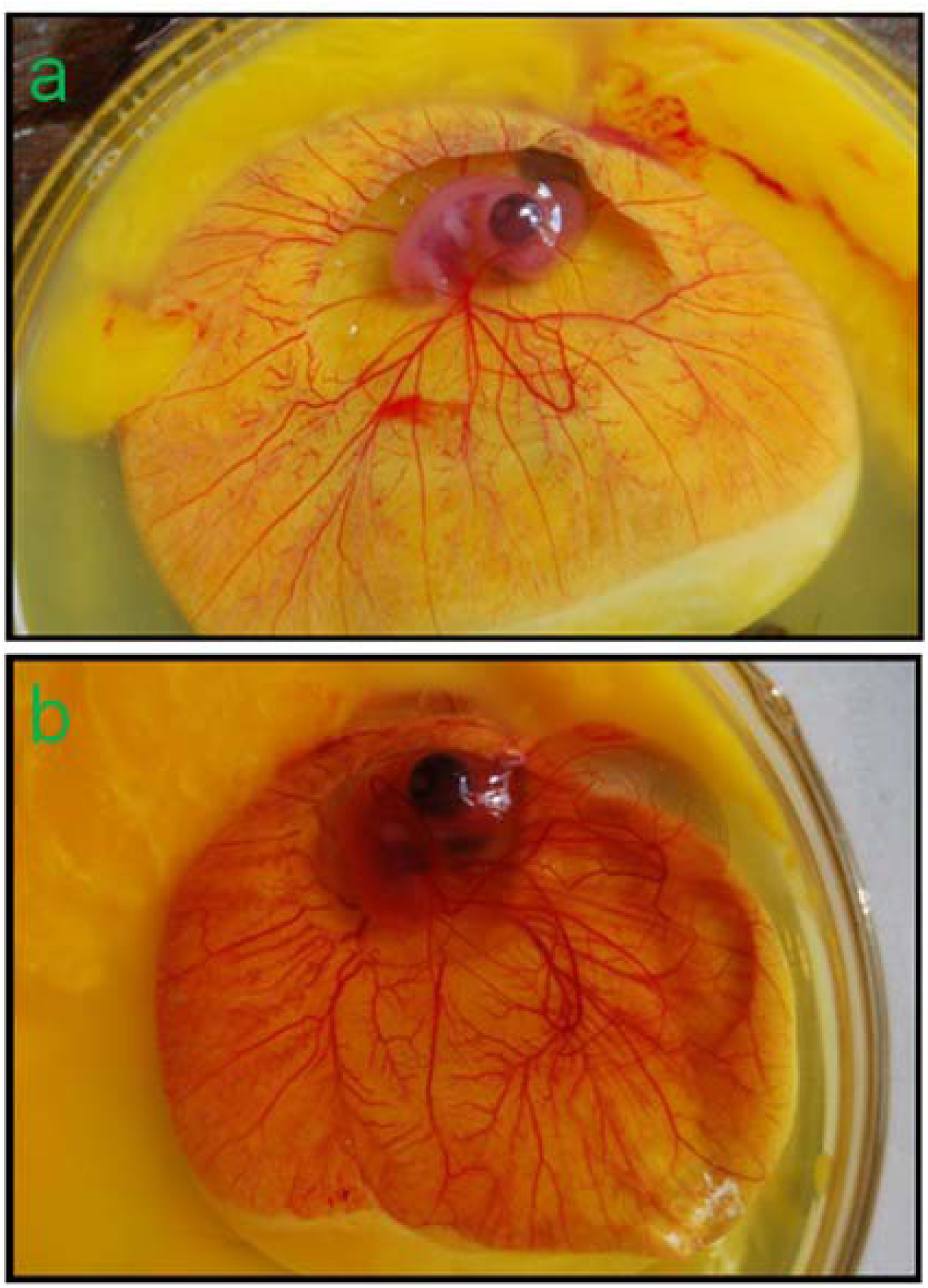
CAM of 9 days old chicken embryo incubated for 5 days (14^th^ day CAM): (**a**) Physiological angiogenesis or control CAM; (**b**) Lysine HCl induced extensive angiogenic response or experimental CAM.

### 3.3. In-silico docking analysis

Z-DOCK result for 4KZN (VEGF-A receptor) and Lysine (as a ligand) shows a specific location on VEGF-A receptor as the most probable binding site for added Lysine (Fig. 2b, c). Interestingly, although negative charge distribution among the entire body of the peptide was uniform (Fig. 2a), added external lysine binding had a preferential binding site (Fig. 2b, c).

Again, as observed in VEGF-A receptor-Lysine complexing study, VEGF-A showed a very dedicated binding site for the added single Lysine moiety, although unlike in VEGF-A receptor, VEGF-A showed few additional distributed comparatively insignificant locations of added Lysine binding sites also (Fig. 2e, f). It is worth noting that the preferential and other lysine binding sites’ distribution in the peptide are all being on either ends or on the tip of projected part of the peptide.

### 3.4. In-silico binding studies of pre-Lysine bound VEGF-A and VEGF-A receptor

Complex-A, i.e. the best obtained structure of lysine bound VEGF-A receptor (4KZN + Lysine, 1:1), binding stability with VEGF-A (1VPF) had been shown and compared based on the global energy changes in interactions between native VEGF-A receptor–VEGF-A (4KZN + 1VPF) and Lysine modified (Complex-A + 1VPF) forms. Change in global energy of binding had been taken as a definite parameter of binding stability; lower the energy more the stability. In Complex-A and VEGF-A (1VPF) binding, energy profile shows more unstable binding (Table 1: Step 1 vs Step 4) compared to the Complex-B, i.e. the best obtained structure of Lysine bound VEGF-A (1VPF + Lysine, 1:1), and VEGF-A receptor (4KZN) binding (Table 1: Step 1 vs Step 5) where further lowering of global energy profiles of the final combine clearly shows an enhanced stability of binding (nearly 40 % lowering indicates a definite shift towards much enhanced stability of the ligand–receptor complex).

**Table 1.** Overall profile of docking experiments.

| Step | Receptor Molecule | Ligand | Comments and/or Results |
| --- | --- | --- | --- |
| 1 | 4KZN (VEGF-A receptor) | 1VPF (VEGF-A) | *Global Energy* = -9.41; Score = 16252 |
| 2 | 4KZN (VEGF-A receptor) | Lysine (in pdb format) | Best structure has been selected and named as Complex-A |
| 3 | 1VPF (VEGF-A) | Lysine (in pdb format) | Best structure has been selected and named as Complex-B |
| 4 | Complex-A | 1VPF (VEGF-A) | *Global Energy* = 2.66; Score =17132 |
| 5 | 4KZN (VEGF-A receptor) | Complex-B | Best result with *Global Energy* = -13.14; Score = 17864 |

Added Lysine (1:1) with VEGF-A and VEGF-A receptor gives more stable complex of the peptide-and-receptor (Fig. 5), thereby inducing enhanced angiogenic response as seen clinically and experimentally (Fig. 3, 4), where added lysine molecule with its functional groups –NH_2_ (2) and –COOH (1) possibly helps in stabilizing “a probable loose-fit” between the ligand and the receptor.

**Fig. 5.**
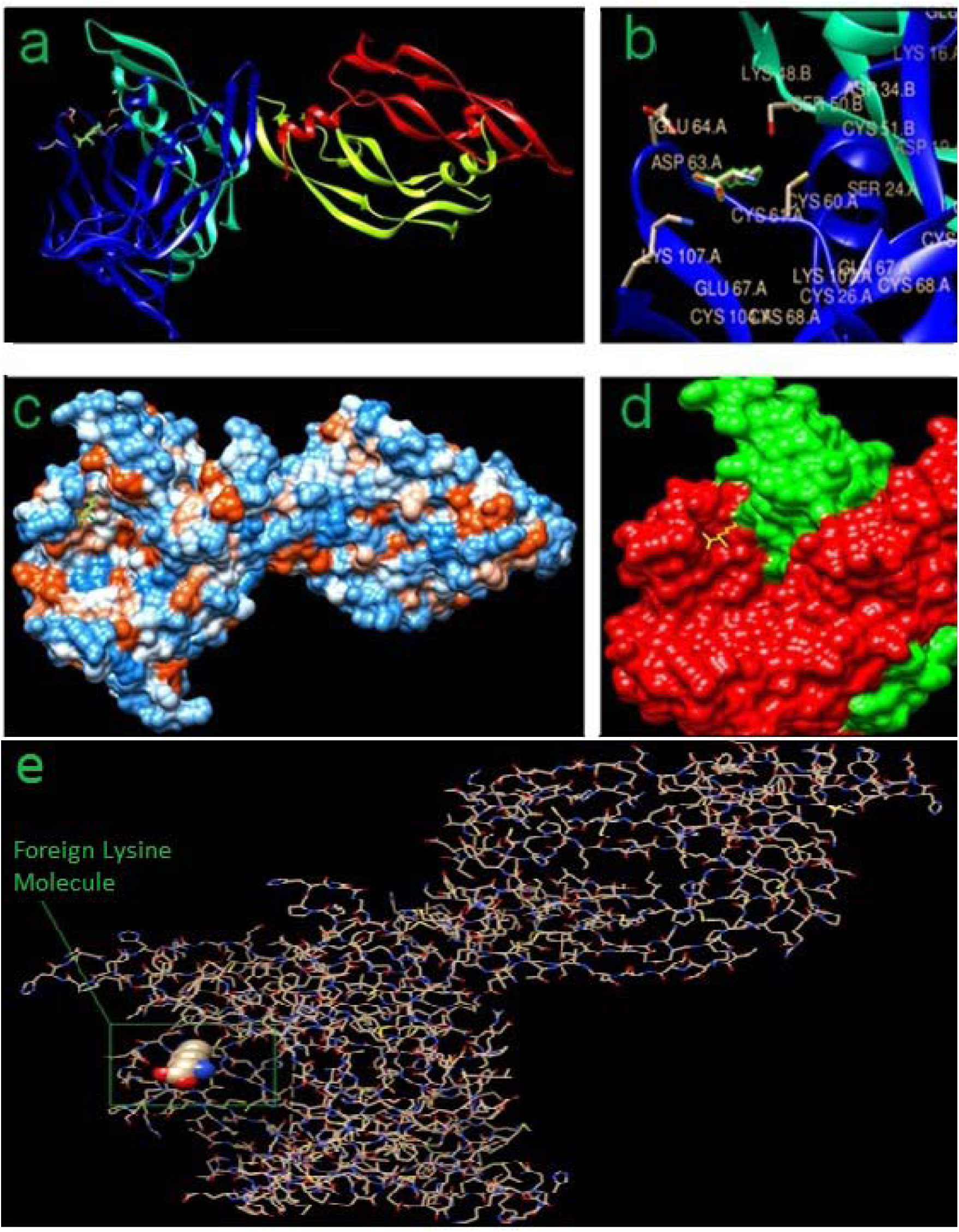
In-silico docking results showing the exact location of “foreign” Lysine molecule: **(a)** Ribbon View; **(b)** Stereo diagram (Foreign Lysine is highlighted with green color); **(c)** Hydrophobic surface view; **(d)** Foreign Lysine in yellow color acting as a binder between VEGF-A (red) and VEGF-A receptor (green); **(e)** All atoms stick view.

## 4. Conclusions

Angiogenesis is a widely conserved process across species which forms the foundations for embryogenesis, organogenesis and all in-situ repair processes like wound repair & tissue regeneration, exchange-bed renewal in all ischaemic diseases like Diabetes and its complications such as Nephropathy, Vasculopathy, Ischaemic Paralytic Stroke, CAD, Placental Insufficiency & IUGR etc. to name a few. Although the process plays such an important role as the foundation of all these divergent conditions, it becomes a fundamentally slow and essentially ineffective process with aging. Embryogenic stage demands an extremely high rate of cell expansion as a basis of organogenesis and overall growth in-utero. The relative anoxic environment may possibly play a key role in the expression level of the key angiogen(s) in foetal tissues and organs. Post birth, this inbuilt stimulus gets removed with possible gradual loss of expression of the peptide and its receptor gene(s) overall on a sustained basis. This possibly explains a very slow collateral generation in ischaemic myocardium with advancing age. A detailed population study would possibly open up answers to differential clinical progresses in patients in ischaemic conditions like Paralytic Stroke, CAD etc., including giving directions to prognostic aspects (objective ischemia assessment) of ongoing therapeutic approaches in all ischemic conditions (e.g. Diabetes and its complications).

Also the level of VEGF-A and VEGFR may well be differentially expressed in various races, communities and even in individuals in a given community and geographical location, based on given genetic profiles. Certain clinical condition like Sudden Cardiac Death may well be explained by near total absence of collateral formation in myocardium with advancing age. Collateral generation and the absence of it can possibly be explained on this molecular expression & interaction status.

Contributing to overall less and differential expression(s) of angiogenic peptide(s) including VEGF-A in individuals, another contributory factor to the “inefficiency” of physiological angiogenic response could be “weak binding” or “nearly no binding” of the peptide and its receptor. The proposed “loose-fit” model may partly explain the overall lack of binding stability leading to the final inefficient biological expression by way of less induction of angiogenic response under normal physiological conditions in ischaemic tissues.

This raises a couple of related questions:

> **1.** Whether titre of VEGF-A (or/and iso-forms) in general circulation or in a particular ischaemic tissue zone gives an idea about the degree of ischaemia? And whether it alone or in combination with a few other Hypoxia induced peptide(s) can indicate definite direction towards objective assessment of a given ischaemic condition, either as an objective diagnostic or prognostic parameter or both. It may as well possibly be a parameter of predictive assessment of ongoing therapy in any of the ischaemic diseases with long drawn complications, like Diabetic Nephropathy, Diabetic Paralytic Stroke, CAD etc. as explained above. This new direction can possibly give advanced indication(s) of setting-in of ischaemia even without any obvious expressions of it or clinical presentations at a given instance (predictive diagnostics).
>
> **2.** Whether a possible new therapeutic angiogenic mode can be visualized by way of the natural amino acid molecule (and its synthetic analogue(s) where in-silico docking work is in progress), despite the molecule per se being non-angiogenic and only possibly helps stabilize the VEGF-VEGFR complex to get its augmented response so long as the two complimentary peptides are in the vicinity in response to an ischaemic stimulus, thereby making the induction process completely controllable.

Basic amino acids (Lysine, Arginine and Histidine) are all vasoactive in addition to their routine metabolic roles. Arginine, in-vivo, acts as the source of NO as one of the most potent vasodilators of existing capillary beds anywhere^29^. Histidine remains locally vasoactive in some tissues - being converted to Histamine.

Vasoactivity of Lysine possibly lies in its ability to form new capillary beds in ischemic tissues and organs as shown in Fig. 3 and 4. Physiological angiogenesis being slow and functionally ineffective in nearly all ischaemic conditions needs augmentation for it to be therapeutically meaningful and a significant mean of controlled reperfusion^30^. Since, it is mediated through VEGF binding to VEGF receptor, the possible proposed “anchorage by the Lysine molecule” enhances the angiogenic response to a very remarkable extent.

Present study raises a possibility of examination of the natural amino acid and its defined synthetic analogue(s) as possible putative agents in clinical induction of time-bound controlled angiogenic responses in ischaemic tissues and organs as an alternate mode of therapeutic approach without any risk of reperfusion injury.

## Acknowledgments

The authors would like to acknowledge the help of Dr. D. Banerjee during the dry experiments.

## Compliance with ethical standards

Clinical studies had been conducted as per the institutional ethical committee guidelines and with informed individual patient consent.

